# Lentiviral CRISPRa/i in the adult Prairie Vole Brain: Modulating Neuronal Gene Expression Without DNA Cleavage

**DOI:** 10.1101/2025.03.30.646142

**Authors:** Meredith K. Loth, Kendall T. Mesch, Celine Herrera-Garcia, Liza E. Brusman, Zoe R. Donaldson

## Abstract

Prairie voles (*Microtus ochrogaster*) are a powerful model for studying the neurobiology of social bonding, yet tools for region- and cell type-specific gene regulation remain underdeveloped in this species. Here, we present a lentivirus-mediated CRISPR activation and interference (CRISPRa/i) platform for somatic gene modulation in the prairie vole brain. This system enables non-mutagenic, titratable regulation of gene expression in the adult brain without germline modification. Our dual-vector system includes one construct expressing dCas9-VPR (CRISPRa) or dCas9-KRAB-MeCP2 (CRISPRi) under a neuron-specific promoter, and a second construct delivering a U6-driven sgRNA alongside an EF1α-driven mCherry reporter. We detail the design, production, and stereotaxic delivery of these tools and demonstrate their application by targeting four genes implicated in social behavior (*Oxtr, Avpr1a, Drd1, Drd2*) across two mesolimbic brain regions: the nucleus accumbens and ventral pallidum. Gene expression analyses confirmed robust, bidirectional transcriptional modulation for select targets, establishing proof of concept for CRISPRa/i in this non-traditional model. The dual-vector design is readily adaptable to other gene targets, cell types, and brain regions, and can be multiplexed to provide a flexible and scalable framework for investigating gene function in behaviorally relevant circuits. These advances represent the first successful implementation of somatic CRISPRa/i in prairie voles and expand the genetic toolkit available for this species.

## 1. Introduction

Bonding and affiliation are fundamental components of social behavior, and their study has implications for understanding both typical and disordered social functioning. Prairie voles (*Microtus ochrogaster*) have emerged as a powerful model species for investigating the neurobiology of social attachment (Sue Carter, Courtney Devries and Getz, 1995; Insel and Young, 2001; Young and Wang, 2004). Unlike traditional laboratory rodents such as mice and rats, but like humans, prairie voles have a naturally monogamous mating system, and exhibit pair bonding behavior and biparental care. This makes them uniquely suited for studying the molecular and neural mechanisms underlying social bonding, partner preference, and parental care. Extensive behavioral characterization, combined with region-specific receptor mapping, have revealed critical roles for oxytocin, vasopressin, and dopamine signaling in the formation and maintenance of social bonds in this species (Young *et al*., 2001; Lim, Murphy and Young, 2004; Aragona *et al*., 2006; Lim and Young, 2006; Donaldson and Young, 2008; Gobrogge and Wang, 2016).

Despite their value as a behavioral model, gene manipulation approaches in voles have historically been more limited compared to other established model organisms. Transgenic prairie voles were first developed over a decade ago (Donaldson *et al*., 2009) and more recent studies have used CRISPR/Cas9 to generate germline knockouts of the oxytocin receptor (Horie *et al*., 2019; Berendzen *et al*., 2023) or to deliver gene-editing tools via AAV to the brain (Boender *et al*., 2023). However, these CRISPR-based approaches have relied on active nuclease strategies, which introduce irreparable genomic damage and are limited to gene disruption.

In contrast, CRISPRa/i systems use catalytically inactive Cas9 (dCas9) fused to transcriptional activators or repressors, enabling potentially reversible, non-mutagenic regulation of endogenous gene expression. While multiple strategies exist for gene manipulation in the brain, few offer the combined advantages of spatial precision, temporal control, and the ability to titrate expression levels in somatic tissues. As summarized in Supplementary Table 1, CRISPRa/i uniquely supports flexible, scalable, and region-specific gene up- or down-regulation without permanent genomic alterations, making it particularly well-suited for probing gene function in behaviorally relevant brain circuits.

The broader application of CRISPRa/i across species is needed to open new directions for comparative neuroscience, behavioral genetics, and evolutionary biology. It will enable targeted, cell-specific manipulation of genes in organisms with ecologically and socially relevant behaviors. Thus, to advance genetic tool development in prairie voles, we established a lentivirus-mediated CRISPRa/i platform for somatic gene regulation in the brain. This system enables activation or interference of target genes in a spatially and cell type-specific manner without requiring germline manipulation. Our protocol involves co-injection of two lentiviral constructs: one encoding the dCas9-VPR (CRISPRa) or dCas9-KRAB-MeCP2 (CRISPRi) effector under the neuron-specific synapsin promoter, and a second carrying a U6-driven sgRNA targeted to the gene of interest with a mCherry reporter under the EF1α promoter. We selected lentivirus for delivery due to its packaging capacity, which accommodates the large size of dCas9-VPR/KRAB-MeCP2 fusion proteins and neuron-specific regulatory elements. In addition, the dual-vector design allows for modular exchange of sgRNA constructs without the need to repackage dCas9, supporting flexible and iterative gene targeting *in vivo*. We validated this approach by modulating expression of oxytocin, vasopressin, and dopamine receptor genes in the nucleus accumbens or ventral pallidum. This platform is adaptable for a range of gene targets, brain regions, and experimental timelines, and holds promise for investigating the molecular basis of social behavior, mapping gene function in specific circuits, and evaluating gene-environment interactions in a behaviorally relevant mammalian model.

## 2. Methods

### 2.1 Animals and Housing

All procedures were approved under the University of Colorado’s Institute of Animal Care and Use Committee (IACUC) and performed in the light phase. All authors complied with the ARRIVE guidelines.

Prairie voles were bred in-house from colonies originating from Emory University and the University of California Davis, with all animals originally descended from wild animals collected in Illinois. Voles were weaned at postnatal day 21 and were then housed in standard static rodent cages (17.5l x 9.0w. x 6.0h. in.) in groups of 2-4 with either same sex siblings or same sex voles from similar weaning time frames. Animals were given *ad libitum* access to water and rabbit chow (5326-3 by PMI Lab Diet). Rabbit chow was supplemented with sunflower seeds, dehydrated fruit bits, and alfalfa cubes. Enriched cages consisted of cotton nestlets, a plastic igloo, and a PVC pipe. Animals were kept in a temperature (23-26 °C) and humidity-controlled room with a 14:10 hour light-dark cycle.

### 2.2 HEK293T Cell Line

HEK293T cells (ATCC, RRID:CVCL_0063) were obtained from the American Type Culture Collection. They were maintained in standard HEK media: DMEM (High Glucose, Pyruvate; Gibco 11995081) supplemented with 10% FBS (Qualified US Origin; BioFluid 200-500-Q) and 1 U/mL Penicillin-Streptomycin (Gibco 15140122). Cells were cultured in T75 or T225 tissue culture flasks and passaged at 70–80% confluence, with a maximum of 25 passages.

### 2.3 Viral Vector Design and Production

Gene-specific sgRNA targets were designed using Benchling (RRID:SCR_013955), the only platform containing the prairie vole genome (MicOch1.0) for identifying potential off-target effects. sgRNA’s were purchased as oligonucleotides with BbsI from Integrated DNA Technologies (IDT). Oligos comprising sgRNAs were annealed together and cloned into the Bbs1 sites in lenti U6-sgRNA/EF1α-mCherry vector (RRID: Addgene_114199). sgRNA’s for CRISPRa were restricted to -500bp upstream of the target gene, and sgRNA’s for CRISPRi were restricted to +300 bp downstream of the target gene as per prior recommendations (Maeder *et al*., 2013; Mali *et al*., 2013; Konermann *et al*., 2015). sgRNAs specificity was assessed using National Center for Biotechnology Information’s (NCBI) Basic Local Alignment Search Tool (BLAST). Information related to sgRNA sequences and distance from gene TSS can be found in Supplementary Table 2. Complete plasmid sequence was confirmed via Plasmidsaurus prior to lentivirus production.

### 2.4 Lentivirus generation

Lentivirus production was performed as specified in Savell et al (Savell, Sultan and Day, 2019) with some modifications. Briefly, large scale viruses were produced in a sterile environment following BSL-2 safety guidelines. HEK-293T cells were transfected with a corresponding CRISPR plasmid: (lenti SYN-FLAG-dCas9-VPR (RRID: Addgene_114196)); lenti SYN-dCas9-KRAB-MeCP2 (RRID: Addgene_155365); lenti U6-sgRNA/EF1a-mCherry (RRID: Addgene_114199)); and psPAX2 packaging plasmid (RRID: Addgene_12260) and the pCMV_VSV-G envelope plasmid (RRID: Addgene_8454) and FuGENE ® HD Transfection Reagent (Promega) in supplemented Ultraculture media (L-glutamine, sodium pyruvate, and sodium bicarbonate) in a T225 culture flask. Supernatant was passed through a 0.45 µm filter and centrifuged at 106,883g (38,100 rpm) for 65 minutes at 4 °C in a 70Ti fixed angle titanium rotor (Beckmann Coulter). The viral pellet was resuspended in 1/100^th^ supernatant volume of sterile PBS and stored at -80°C. Physical viral titer was determined using Lenti-X qRT-PCR Titration kit (Takara) and only viruses with > 1 × 10^9^ GC/ml were used. Viruses were stored in sterile aliquots of PBS at -80°C.

### 2.5 Stereotactic surgeries and viral injection protocol

Naïve adult prairie voles were anesthetized with isoflurane (4% induction and 1.5-2.5% maintenance) at an oxygen flow rate of 1L/min and secured in a head-fixed stereotaxic frame (Kopf Instruments). Body temperature was maintained at 37°C using a closed loop heating pad with a rectal thermometer. Eyes were lubricated with ophthalmic ointment (Sterile Lubricant Eye Ointment), and depth of anesthesia was monitored by breathing and toe and tail pinch response. Using a shaver, fur was removed from the dorsal portion of the head and, a midline incision was made and disinfected with 70% isopropyl alcohol and betadine. Briefly, the scalp and connective tissue were removed above the frontal skull plates and the head leveled in the anterior-posterior plane. Guide holes were drilled using stereotaxic coordinates (all coordinates in respect to bregma (Paxinos and Watson, 2007): nucleus accumbens: AP: +1.7; ML ±: 1.0; ventral pallidum: AP: +1.5; ML ±: 0.9; All infusions were made using a gastight 30-gauge stainless steel injection needle (Hamilton Syringes) that extended into the infusion site. Bilateral lentivirus microinfusions were made using a UMP3-T syringe pump (World Precision Instruments) at a rate of 2nL/sec at DV: - 4.8/4.7/4.6 mm for the nucleus accumbens or DV: -5.8/5.7/5.6 mm for the ventral pallidum. At each DV coordinate, 700/700/600 nl were injected for a total of 2 uL per hemisphere (See Supplemental Fig 1 for virus volume validation). Injection needles remained in place for 10 minutes post infusion to allow time for diffusion. Prairie voles were infused bilaterally with 2 uL of total lentivirus mix which comprised of a 1:10 ratio of sgRNA virus to dCas9-VPR or KRAB-MeCP2 virus in sterile PBS. Surgical incisions sites were closed with vicryl sutures. Animals received extended release meloxicam (4mg/kg), lidocaine at the incision site, and saline (1 mL) for pain, infection prevention, and post-operative management. Animals recovered 72 hours in a BSL-2 facility to minimize any exposure to animal shedding of lentivirus particles.

### 2.6 Tissue collection and microdissection

For samples used in qRT-PCR, animals were euthanized via rapid decapitation. Brains were immediately extracted and rinsed in sterile saline. Fluorescent guided dissections were performed manually using sterile RNase free razor blades and mCherry fluorescence was visualized with a fluorescent dissecting microscope Olympus MVX10 MacroZoom at the University of Colorado Boulder MCDB Light Microscopy Core Facility (RRID:SCR_018993). During dissection, tissue was kept on ice blocks and in cold saline. Dissected hemispheres were flash frozen on dry ice and stored at -80°C.

For samples used for immunohistochemistry, animals were injected with 0.15-0.30 mL 1:2 ketamine/xylazine and transcardially perfused with 4% paraformaldehyde (PFA, Electron Microscopy Scienes) in phosphate buffered saline. Brains were kept in 4% PFA overnight and transferred to 30% sucrose. After brains sank, they were rinsed and flash frozen on dry ice and stored at -80°C for 24 hours prior to sectioning on a freezing microtome (Leica June SM2000R, 40 µm/slice)

### 2.7 RNA Extraction and cDNA generation

Samples were processed for total purified RNA as described in Cunningham et al (Cunningham *et al*., 2019) using the Norgen Total RNA/gDNA kit (Norgen Biotek no. 48700). Briefly, frozen tissue was placed in 600 µL ice cold lysis buffer and homogenized using a Scilogex homogenizer. Homogenized tissue was kept on wet ice while all animals for the cohort were processed. The homogenizer tip was thoroughly cleaned in between each animal with DEPC-treated water and 70% ethanol to prevent cross contamination across samples. Homogenized samples were centrifuged at 4000 x g for 10 minutes and supernatant was transferred to new pre-chilled tubes.

Total RNA (2 uL for each sample) was Nanodropped to determine RNA concentration (Agilent Technologies). RNA samples were all diluted to 25 ng/µl with RNAse-free water (Norgen). cDNA was generated using high Capacity cDNA Reverse Transcription kit (Applied Bio Systems). For each sample, cDNA was produced using a mixture of 2 μL of 10X RT buffer, 0.8 μL of dNTP, 2 μL of random primers, 1 μL of Transcriptase, 1 μL of RNA inhibitor, and 3.2 μL of ddH2O. RNA (250 ng) was then added, and cDNA was generated using reaction conditions 25°C for 10 min, 2 × (37 °C for 120 min, 85 °C for 5 min). Samples were stored at −20 °C until processing.

### 2.8 qRT-PCR validation of gene expression

Prairie vole gene expression for *Oxtr, Avpr1a, Drd1, Drd2*, and *Gapdh* was quantified using qPCR. Primer sequences are provided in Supplementary Table 3. Samples and probes were processed in triplicate in MicroAmp Fast Optical 96 well Reaction Plate (Applied Bio Systems) with 0.66 µl of cDNA, 0.50 µl of probe, 5 µl of TaqMan Fast Advanced Master Mix (Applied Biosystems), and 3.84 µl of ddH_2_O per well for a total volume of 10 µl. Plates were covered with Optical Adhesive Film (Applied Biosystems), vortexed, and centrifuged prior to PCR amplification in Applied Biosystems QuantSTudio 3 qPCR machine (Applied Biosystems). DNA was amplified using the following cycling conditions: 50 °C for 2 min, 95°C for 20 s, and 40 cycles of 95°C for 1 s and 60 °C for 20 s.

The mean cycle threshold (Ct) value was calculated for each sample run in triplicate. Triplicates with a standard deviation greater than 0.30 were assessed for potential outliers. If no clear outlier was visually identified, the sample was excluded from analysis. Gene expression quantification was performed using the ΔΔCt method (Livak and Schmittgen, 2001), with all samples normalized to *Gapdh*.

### 2.9 Immunohistochemistry and Microscopy

Immunohistochemistry was performed to visualize the FLAG tag on dCas9-VPR and dCas9-KRAB-MeCP2 constructs. Free-floating brain slices from PFA-perfused prairie voles were stored at -20°C in cryoprotectant consisting of 50% glycerol in 0.05 M phosphate buffer. For immunofluorescent labeling of FLAG, sections were washed in 1X PBS, for 30 minutes, with fresh PBS every 5 min. Sections were blocked with 5% normal donkey serum (Jackson Laboratories) containing 0.2% Triton X-100 for 1 hr followed by primary antibody incubation with mouse-anti-FLAG (Invitrogen, RRID:AB_1957945, 1:100) at 4°C for 40 hrs in 750 uL of solution. After washing, sections were incubated with AlexaFluor 488 secondary antibody (Life Technologies, 1:500) for 1 hr. Slices underwent another series of washes and were then mounted on slides and coverslipped with Prolong Gold Antifade Mountant (Life Technologies) to preserve signal intensity and brightness.

Slices were imaged using a Nikon A1 Laser Scanning confocal Microscope at the University of Colorado Boulder MCDB Light Microscopy Core Facility (RRID:SCR_018993). Confocal images were taken with a 20x lens in two channels (green and red) of the nucleus accumbens and ventral pallidum. Confocal stacks were projected as single images using maximum fluorescence.

### 2.10 Data analysis and Statistics

Data are shown as means ± standard error of the mean. Statistical significance α was set at 0.05. All n values represent number of animals. Statistical analyses were carried out using either GraphPad PRISM (version 10.4.1, Graphpad, San Diego, CA) or Python (v 3.12.4) in a reproducible computing environment. Python scripts were executed in Juypter Notebook (v 7.0.8) via Anaconda Navigator (v 2.6.4). Paired t-tests with error propagation were performed.

## 3. Results

### 3.1 CRISPR a/i constructs successfully expressed in the prairie vole brain

We employed a dual-vector system to enable gene activation or repression. One lentiviral construct encoded either dCas9-VPR (Fig. 1A) or dCas9-KRAB-MeCP2 (Fig. 1B) under the control of the neuron-specific SYN promoter. The second construct carried a U6-driven sgRNA targeting either *Oxtr, Avpr1a, Drd1, Drd2*, or a non-targeting *lacZ* control, and included a mCherry reporter driven by the EF1α promoter for visualization of sgRNA-expressing cells (Fig 1A-B). Following lentiviral injection, animals were given 21 days to allow for both recovery and robust expression of CRISPRa/i constructs (Fig. 1C), after which immunohistochemistry (IHC) was performed to confirm expression in targeted brain regions, the nucleus accumbens (Fig 1D) or ventral pallidum (Fig 1E).

**Figure 1.**
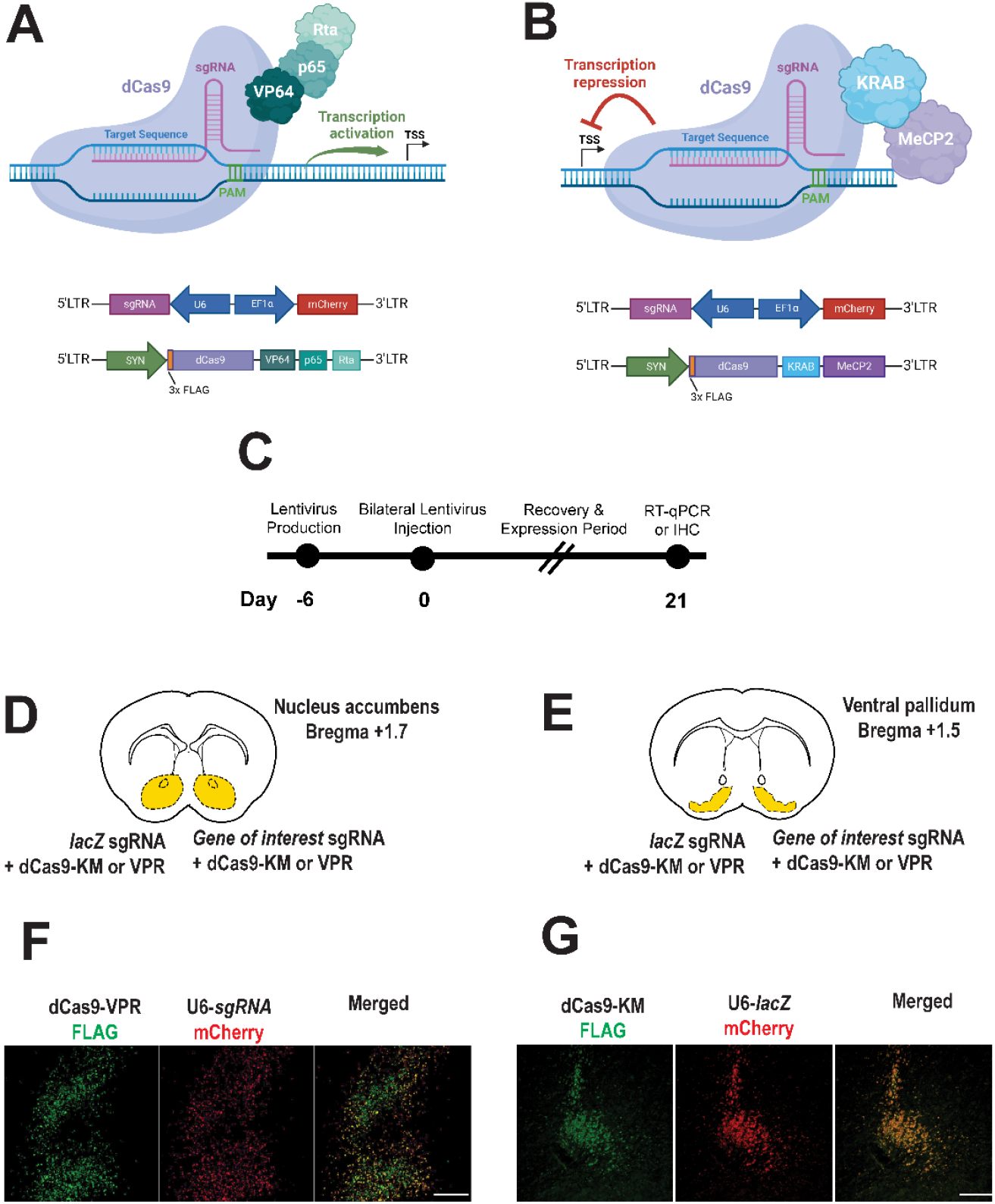
CRISPR a/i constructs successfully expressed in the prairie vole brain. **A-B**. CRISPRa vector for expressing dCas9-VPR activator fusion (**A**), CRISPRi vector for expressing dCas9-KRAM-MeCP2 interference fusion (**B**), and sgRNA’s targeting either the bacterial *lacZ* gene (nontargeting control) or other genes of interest in the prairie vole (**A-B**). **C**. Experimental timeline for CRISPRa or CRISPRi expression the prairie vole brain (nucleus accumbens or ventral pallidum). **D-E**. Brain atlas diagrams of the nucleus accumbens (**D**) and the ventral pallidum (**E**) indicating dual bilateral injections of CRISPRa or CRISPi construct with sgRNA construct targeting lacZ(control) or gene of interest. **F-G**. Immunohistochemistry images reveal successful transduction and show representative expression of CRISPRa (**E**) and CRISPRi (**F**) (FLAG, green) lentiviruses along with co-expression of U6-driven sgRNA’s (mCherry, red). Merged images confirm colocalization of CRISPRa/i with sgRNA viruses (scale bar = 100 µm).

Representative images demonstrate detection of constructs in the nucleus accumbens (1F) and the ventral pallidum (Fig 1G). Specifically, FLAG-tagged dCas9-VPR (CRISPRa) and FLAG-tagged dCas9-KM (CRISPRi) were detected in transduced neurons (Fig. 1F and G, left panels, respectively). Co-expression of U6-driven sgRNA’s, co-labeled with a mCherry reporter under an EF1α, promoter was observed in the same regions (Fig. 1F and G, middle panels). Merged images confirm colocalization of dCas9 constructs and sgRNA’s indicating successful neuronal expression of both components (Fig. 1F and G, right panels).

These findings validate expression of lentiviral CRISPRa/i in a non-traditional mammalian model, demonstrating effective lentiviral-mediated delivery and neuronal expression of CRISPRa and CRISPRi constructs in the prairie vole brain across multiple brain regions.

### 3.2 CRISPR a/i modulation of behaviorally relevant genes

To evaluate the effectiveness of CRISPRa and CRISPRi in the prairie vole brain, we focused on four target genes implicated in social behavior: the nonapeptide receptors *Oxtr* and *Avpr1a*, and the two most abundant dopamine receptors, *Drd1* and *Drd2*. We targeted these genes in mesolimbic brain regions where they are highly expressed and known to modulate pair bonding behaviors (Wang *et al*., 1999; Young, 1999; Gingrich *et al*., 2000; Young *et al*., 2001; Aragona *et al*., 2006; Gobrogge and Wang, 2016; Loth and Donaldson, 2021; Pierce *et al*., 2024). Specifically, *Oxtr, Drd1*, and *Drd2* were targeted in the nucleus accumbens (NAc), while *Avpr1a* was targeted in the ventral pallidum (VP).

#### 3.2.1 Oxtr mRNA Expression

qPCR analysis revealed that CRISPRa significantly increased *Oxtr* expression, while CRISPRi led to a significant reduction (Fig. 2 A and B, respectively). Specifically, dCas9-VPR activation resulted in a 4.29-fold increase in *Oxtr* expression compared to lacZ control (paired t-test, p = 0.03). Conversely, dCas9-KRAB-MeCP2 repression significantly decreased *Oxtr* expression by 87% (p < 0.001). These findings confirm that CRISPRa/i tools can effectively regulate *Oxtr* transcription in the prairie vole brain.

**Figure 2.**
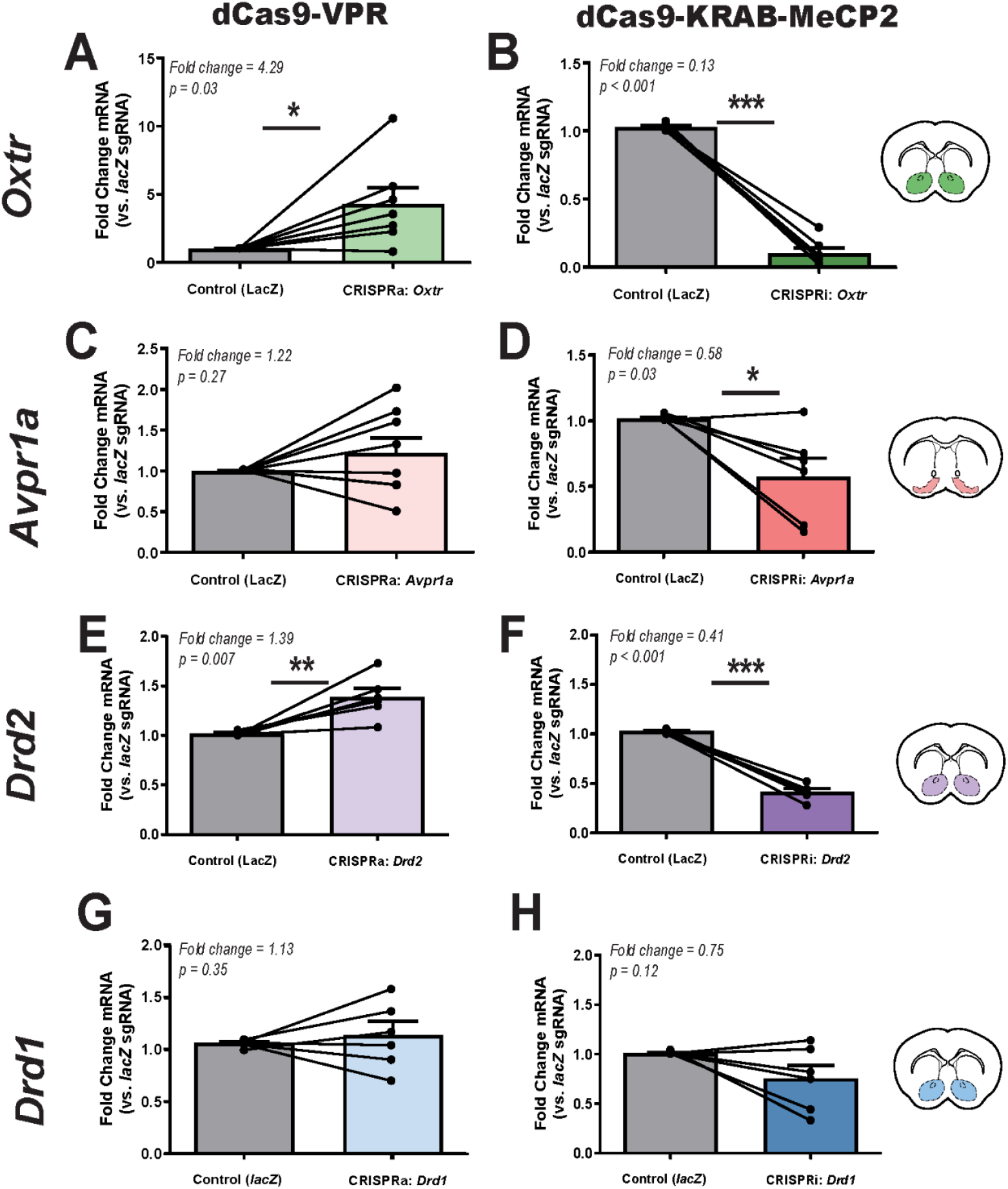
CRISPRa and CRISPRi modulation of target gene expression in the prairie vole brain. **A-B**. Relative *Oxtr* mRNA expression following CRISPRa (dCas9-VPR, **A**) and CRISPRi (dCas9-KRAB-MeCP2, **B**) manipulation in the nucleus accumbens, as measured by qPCR. **C-D**. Relative *Avpr1a* mRNA expression following CRISPRa (**C**) and CRISPRi (**D**) in the ventral pallidum. **E-F**. Relative *Drd2* mRNA expression following CRISPRa (**E**) and CRISPRi (**F**) in the nucleus accumbens. **G-H**. Relative *Drd1* mRNA expression following CRISPRa (**G**) and CRISPRi (**H**) in the nucleus accumbens Injection sites are illustrated in coronal sections (inset), with colored markers indicating targeting of the nucleus accumbens (green, purple, blue) and ventral pallidum (pink). Each animal served as its own internal control, with *lacZ* control sgRNA injected into the left hemisphere and gene-specific sgRNA injected into the right hemisphere. Expression levels were normalized to *Gapdh*, and data are presented as fold change relative to l*acZ* control (mean ± SEM). Statistical significance was determined using paired t-tests, with p < 0.05 considered significant (*, p < 0.05; ***, p < 0.001).

#### 3.2.2 Avpr1a mRNA Expression

qPCR analysis revealed greater variability in Avpr1a expression following CRISPRa and CRISPRi manipulation (Fig 2C-D). CRISPRa-driven activation led to a 1.22-fold increase in *Avpr1a* expression relative to the control, but this change was not statistically significant (*p* = 0.27). In contrast, CRISPRi significantly reduced expression by 42% (*p* = 0.03). Notably, *Avpr1a* expression levels exhibited high variability across animals and the magnitude of these effects were smaller than those observed for *Oxtr*. An additional CRISRPi sgRNA tested for *Avpr1a* yielded similar results with significantly reduced expression by 37% (p = 0.02) (Supplemental Fig. 2). However, the overall trend of CRISPRa upregulation and CRISPRi repression was consistent.

#### 3.2.3 Drd2 mRNA Expression

Drd2 expression was modulated in a gene-specific manner, with CRISPRa inducing a significant 1.39-fold increase (p = 0.007) and CRISPRi leading to a 59% reduction (p < 0.001) (Fig.2G-H). Variability in *Drd2* expression across animals was lower than *Drd1* and *Avpr1a*, which may have contributed to its statistical significance. While Drd2 expression was significantly modulated by CRISPRa and CRISPRi, the magnitude of these changes was smaller compared to *Oxtr*, suggesting potential differences in baseline expression levels or regulatory mechanisms within the nucleus accumbens.

#### 3.2.4 Drd1 mRNA Expression

Similar to *Avpr1a, Drd1* expression exhibited variability across animals following both CRISPRa and CRISPRi manipulation (Fig 2 E-F). CRISPRa injection resulted in a 1.13-fold increase in *Drd1* expression, though this change was not statistically significant (p = 0.35). Similarly, CRISPRi injection led to a 25% reduction in *Drd1* expression, but this effect was also not significant (p = 0.12).

Taken together, these results confirm the feasibility of using CRISPRa/i to modulate *Oxtr* and *Drd2* expression in prairie voles, while highlighting potential technical limitations in targeting *Avpr1a* and *Drd1* with the current approach.

## 4. Discussion

Our findings establish lentivirus-mediated CRISPRa/i as a flexible and regionally precise tool for manipulating somatic gene expression in prairie voles. By delivering lentiviral constructs encoding dCas9-VPR or dCas9-KRAB-MeCP2 alongside U6-driven sgRNAs, we demonstrate the feasibility of bidirectional transcriptional control in neurons of the adult prairie vole brain. This system provides a valuable method for probing gene function *in vivo*—particularly for genes implicated in complex social behaviors—without requiring germline genetic manipulation. While *in vivo* CRISPRa/i systems have been validated in mice and rats (Savell *et al*., 2019, 2020; Duke *et al*., 2020; Deng *et al*., 2022; Bendixen, Jensen and Bak, 2023), to our knowledge, this is the first application in a non-traditional mammalian model. Expanding gene modulation tools to prairie voles opens new opportunities to study the molecular basis of social bonding in a species with affiliative behaviors more analogous to humans.

We demonstrate effective region-specific modulation of key social behavior genes across two distinct brain regions: the nucleus accumbens (*Oxtr, Drd1, Drd2*) and the ventral pallidum (*Avpr1a*). *Oxtr* and *Drd2* showed robust, consistent modulation following injection into the nucleus accumbens, confirming effective somatic gene regulation using CRISPRa/i. These results provide proof-of-concept for somatic, region-specific gene activation and interference in prairie voles. The ability to efficiently upregulate or downregulate key neuromodulatory receptors—without permanent genomic alterations—offers a powerful platform for studying the dynamics of oxytocinergic and dopaminergic signaling in behaviors ranging from pair bonding and biparental care.

However, not all targets exhibit equal efficiency of repression or activation. *Avpr1a* and *Drd1* exhibited more variable modulation, highlighting areas for continued methodological refinement. For *Avpr1a*, CRISPRi led to a modest but statistically significant reduction in expression, while CRISPRa effects were inconsistent and did not reach statistical significance. The two CRISPRi sgRNAs, positioned at +115 and +257 bp relative to the transcription start site (TSS), yielded comparable outcomes, suggesting that TSS proximity alone does not explain the variability. Other factors—such as chromatin accessibility, local epigenetic context, or sequence-specific properties—may have influenced sgRNA efficacy at this locus. The presence of an *Avpr1a* pseudogene in prairie voles did not impact interpretation, as our qPCR primers were designed to exclude pseudogene amplification and the gRNAs targeted sequences specific to the functional gene (Young *et al*., 1997). Variability in *Drd1* modulation may also reflect a combination of locus-specific and technical factors, such as lentiviral transduction variability, promoter interference, or subject-level biological differences. Additionally, sequence variation at or near sgRNA binding sites, such as naturally occurring SNPs (single nucleotide polymorphisms), could impact targeting efficiency in outbred populations and warrants future investigation. These findings highlight the need for systematic sgRNA validation and dose-response/multiplex testing as part of CRISPRa/i implementation in novel species.

Despite these gene-specific challenges, CRISPRa/i offers multiple advantages over nuclease-based editing. Because dCas9 lacks catalytic activity, this system avoids introducing DNA double-strand breaks, thereby preserving genomic integrity and enabling reversible gene regulation. This is particularly advantageous for somatic studies in the brain, where permanent edits may trigger developmental compensation or long-term side effects. In addition, CRISPRa/i allows for precise spatial and temporal control—gene expression can be manipulated after development, in specific brain regions, and within defined cell populations without altering the germline or affecting non-target tissues. CRISPRa/i also enables tunable gene expression, allowing researchers to modulate transcriptional output rather than merely switching genes on or off. Finally, the system supports multiplexed sgRNA delivery for simultaneous regulation of multiple genes within a circuit. While not the focus of the present study, these capabilities underscore the broader utility of the CRISPRa/i platform for dissecting complex gene networks and behaviorally relevant pathways.

Another key strength of CRISPRa/i is its gene-level specificity. Unlike pharmacological agents, which often target multiple receptor subtypes within a class (e.g., D1-like or D2-like dopamine receptors), sgRNAs can be designed to selectively target a single gene. This specificity is particularly valuable when dissecting the roles of closely related receptor isoforms in complex behaviors. For example, while commonly used D2 receptor antagonists can also bind D3 receptors (Scatton *et al*., 2001; Bock *et al*., 2004; Stahl, 2017), CRISPRi targeting of *Drd2* allows for specific repression of *Drd2* transcription without affecting *Drd3*. Similarly, pharmacological tools for oxytocin and vasopressin receptor systems frequently exhibit cross-reactivity due to high sequence homology, whereas CRISPRa/i enables selective modulation of *Oxtr* or *Avpr1a* independently. This specificity is crucial for resolving gene-specific contributions to behavior in systems where receptor subtypes have overlapping yet distinct roles.

There are a handful of limitations to our advance. While qPCR confirmed effective transcriptional modulation, we were unable to validate changes at the protein level due to limited antibody availability for our targets. Although autoradiography could potentially be used for receptor-level detection, it was not implemented here due to resource constraints. Additionally, we did not assess behavioral outcomes, and thus the functional impact of gene modulation remains to be tested in future work. Lentiviral vectors require BSL-2 containment and may not be optimal for all applications. Lastly, the long-term stability of lentiviral CRISPRa/i expression and the possibility of off-target effects warrant further investigation.

Despite these challenges, this work provides a critical proof-of-concept for somatic gene regulation using CRISPRa/i in prairie voles. Beyond validating a powerful new method, our results highlight key opportunities for future development:

- Refining sgRNA selection pipelines, particularly for AVPR1A inhibition, including exploration of sequence variation at sgRNA target sites as a potential contributor to guide efficiency
- Exploring alternative viral platforms (e.g., AAV) using smaller dCas9 variants
- Expanding this system to non-neuronal cell types via other cell-type specific promoters
- Enhancing temporal precision through inducible dCas systems
- Investigating potential off-target activity or compensatory gene expression *in vivo*
- Assessing behavioral consequences of gene modulation in social bonding paradigms

In summary, we establish lentivirus-mediated CRISPRa/i as a novel and effective method for somatic gene regulation in prairie voles, capable of modulating gene expression in a region- and cell type-specific manner across multiple behaviorally relevant brain regions. This system fills a critical methodological gap for genetic manipulation in this species, laying the foundation for future work on the molecular mechanisms of social behavior.

## Author contributions

MKL: Conceptualization, Data curation, Formal analysis, Funding acquisition, Investigation, Methodology, Software, Supervision, Visualization, Writing-original draft, Writing-review & editing. KTM: Formal analysis, Investigation, Writing-review &editing. CHG: Formal analysis, Investigation, Writing-review & editing. LEB: Investigation, Methodology, Writing-review & editing. ZRD: Conceptualization, Funding acquisition, Methodology, Project administration, Resources, Supervision, Writing-original draft, Writing-review & editing.

## Acknowledgements

We thank the voles for their sacrifice and contribution to research. We thank Jessica Abazaris and the rest of the animal care staff at the University of Colorado Boulder for their excellent care of the voles. Kelly Winther, Katie Gallagher, and Kresil Gordon managed the animal colony and provided experimental support. We acknowledge the Light Microscopy Core Facility, Porter B047, B049, B051 and B059 at the University of Colorado Boulder (RRID:SCR_018993) for help and advice with microscopy and thank Dr. James D. Orth for his assistance. We thank the Donaldson lab, Devanand Manoli’s lab, and Jessica Tollkuhn’s lab for their advice, feedback and support. This work was supported by awards from the Dana Foundation, the Whitehall Foundation, National Science Foundation (NSF) IOS-1827790, and National Institute of Health (NIH) DP2OD026143 to Z.R.D. and T32 DA 17637 support to M.K.L.

## Data Availability Statement

Datasets are available on request. The raw data supporting the conclusions of this article will be made available by the authors, without undue reservation.

