## Supplementary Materials for "Lentiviral CRISPRa/i in the adult Prairie Vole Brain: Modulating Neuronal Gene Expression Without DNA Cleavage"

### *Supplementary Material*

1. Supplementary Figures and Tables
   1.1. Supplementary Figures

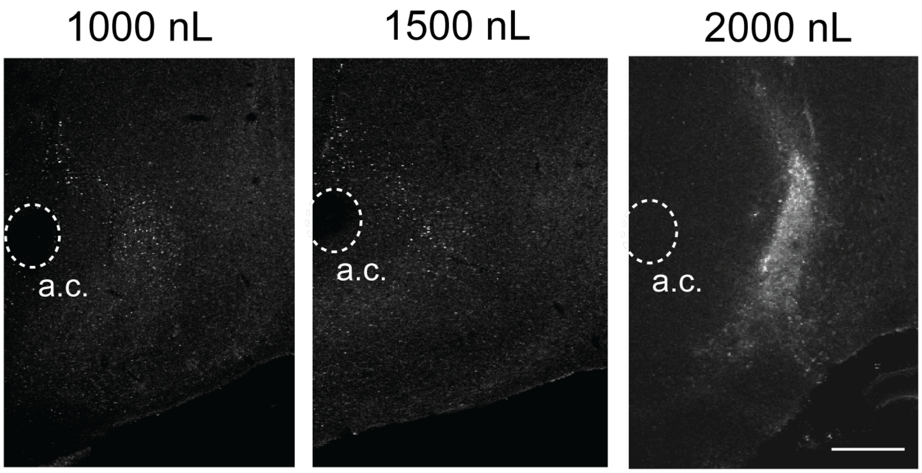


**Supplementary Figure 1.** Virus Volume Optimization for stereotaxic infusion. Representative images of mCherry fluorescence in the prairie vole brain following stereotaxic infusion of increasing viral volumes (1000 nL, 1500 nL, and 2000 nL). Images illustrate spread and expression intensity of lentiviral constructs in the nucleus accumbens, with higher volumes producing greater diffusion and signal intensity. These data informed selection of 2000 nL as the optimal infusion volume for subsequent experiments. Scale bar = 500 µm. a.c. = anterior commissure

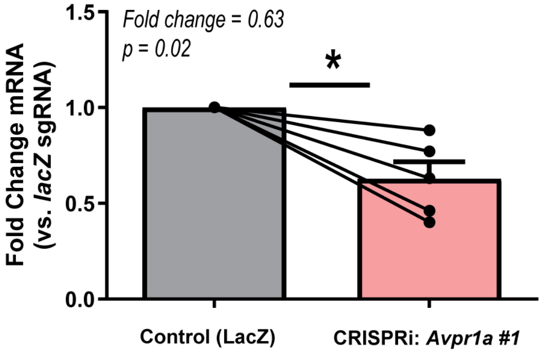


**Supplementary Figure 2.** Validation of an additional CRISPRi guide RNA targeting *Avpr1a*.
qPCR analysis of *Avpr1a* expression in the ventral pallidum following CRISPR interference using an alternative sgRNA (*Avpr1a* #1) compared to a non-targeting control (LacZ sgRNA). Each line represents a paired hemisphere from a single animal. Expression was normalized to *Gapdh* and presented as fold change relative to *lacZ* control. This guide yielded a modest but significant reduction in *Avpr1a* expression, consistent with the primary sgRNA used in the main text.

#### 1.2. Supplementary Tables

**Supplementary Table 1.** Comparison of genetic manipulation methods across key functional criteria.

| Method | Non-mutagenic | Multiplexing capable | Manipulates endogenous expression | Same system for up & down regulation | Titratable gene expression | Potentially reversible |
|---|---|---|---|---|---|---|
| **CRISPRa/i** *(this study)* | ✓ | ✓ | ✓ | ✓ | ✓ | ✓ |
| AAV-mediated overexpression | ✓ | ✓ | ✗ | ✗ | ✓ | ✓ |
| Germline CRISPR NHEJ | ✗ | ✗ | ✓ | ✗ | ✗ | ✗ |
| Germline CRISPR HDR | ✗ | ✓ | ✓ | ✗ | ✗ | ✗ |
| AAV Cas9 KO | ✗ | ✓ | ✓ | ✗ | ✗ | ✗ |
| RNAi | ✓ | ✓ | ✓ | ✗ | ✓ | ✓ |

This table compares six genetic manipulation strategies based on their ability to achieve core experimental features: non-mutagenic manipulation, multiplexing capability, modulation of endogenous gene expression, use of the same system for both up- and down-regulation, titratable expression, and potential reversibility. CRISPRa/i (this study) supports all six features, highlighting its flexibility for somatic gene regulation in the brain.

**Abbreviations:** CRISPRa/i, CRISPR activation/interference; AAV, adeno-associated virus; CRISPR NHEJ, CRISPR-mediated non-homologous end joining; CRISPR HDR, CRISPR-mediated homology-directed repair; KO, knockout; RNAi, RNA interference.

**Supplementary Table 2.** sgRNA sequences and position relative to transcription start site (TSS) for CRISPRa/i targeting in prairie voles

| Target gene (Prairie Vole) | CRISPR Modality | sgRNA sequence (5' → 3') | Position Relative to TSS |
|---|---|---|---|
| *Oxtr* | CRISPRa | GCGGCGACACTCCACTCCCG | -60 |
| | CRISPRi | GCCTGAAACAAACCGAGAGG | +130 |
| *Avpr1a* | CRISPRa | GGGGACTGGAGCAAACCCTG | -102 |
| | CRISPRi | GTTAGGACAGGCTTTCTCGG | +257 |
| *Drd2* | CRISPRa | GCAGTGTAGAGATTGCACTC | -113 |
| | CRISPRi | GCGCGGGGAACCAGGAGCGG | +48 |
| *Drd1* | CRISPRa | GGAATAACATGCTAGCCGAA | -286 |
| | CRISPRi | GGGGTGCTTACCGCTCCGGG | +95 |

Each row lists the target gene (*Oxtr*, *Avpr1a, Drd2, Drd1*), CRISPR modality (CRISPRa = activation, CRISPRi = interference), sgRNA sequence (5′–3′), and its position relative to the transcription start site (TSS). sgRNAs were designed using Benchling (RRID:SCR_013955) to maximize predicted on-target efficiency and minimize off-target effects, based on scoring algorithms from Doench et al. (2016). Scores were generated using the guide-sequence–only model due to incomplete gene annotations in the prairie vole genome. Genomic coordinates and positions relative to the transcription start site (TSS) were derived from the MicOch1.0 genome assembly or updated annotations provided by collaborators. Sequences were selected within ±300 bp of the TSS, with upstream regions prioritized for CRISPRa and downstream regions for CRISPRi.

**Abbreviations:** CRISPRa = CRISPR activation (dCas9-VPR); CRISPRi = CRISPR interference (dCas9-KRAB-MeCP2); TSS = transcription start site.

**Supplementary Table 3.** qPCR Primer and Probe Sequences

| Target gene (Prairie Vole) | Primer Sequences | Probe |
|---|---|---|
| *Oxtr* | 5'-TTCGTGCAGATGTGGAGC-3' | 5'-/56-FAM/CAACCCCTG/ZEN/ GATCTACATGCTGTTCA/31ABkFQ/-3' |
| | 5'-AGTTCGTGAAAGAGGTGGC-3' | |
| *Avpr1a* | 5'-GGGAAATAGTCTTCACGCTGCTGACA-3' | 5'-/56-FAM/TTGTGGAAG/ZEN/ GGAGCCACGTCTTC/31ABkFQ/-3' |
| | 5'-GCTATGGCTTCATCTGCTACCACATCT-3' | |
| *Drd1* | 5'-CATACGTCCTGCTCAACCTG-3' | 5'-/56-FAM/TTGGCATCC/ZEN/ TCGGTGTCTTCCAG/31ABkFQ/-3' |
| | 5'-CTCATCTCCTTTATCCCAGTGC-3' | |
| *Drd2* | 5'-GACCTCCCTTAAGTCAATGAGC-3' | 5'-/56-FAM/TGTGTGGCT/ZEN/ TTCTTCTCCTTCTGCTG/31ABkFQ/-3' |
| | 5'-AGTGTATGTTCAGGATGTGCG-3' | |
| *Gapdh* | 5'-CCACTCTTCCACCTTCGATG-3' | 5'-/56-FAM/TTGAGAGCA/ZEN/ ATGCCAGCCCCA/31ABkFQ/-3' |
| | 5'-GCCAAATTCATTGTCGTACCAG-3' | |

Primer and probe sequences used for gene expression analysis of target genes in prairie vole (*Microtus ochrogaster*) brain tissue. Primers are listed in 5'–3' orientation. All probes are dual-labeled with 5′ 6-FAM fluorophore, internal ZEN quencher, and a 3′ Iowa Black FQ (31AbkFQ) quencher.

*Oxtr, Avpr1a, Drd1, Drd2* = target genes; *Gapdh* = endogenous control gene.
